# The relationships among alkaloid production, fungal biomass, and host genetics in the Tall Fescue-Epichloë symbiosis

**DOI:** 10.1101/2025.05.30.657052

**Authors:** Darrian R. Talamantes, Courtney Phillips, Carolyn Young, Jason G. Wallace

## Abstract

Tall fescue (*Lolium arundinaceum*) is an important forage and turf grass that covers 35 million acres (140,000 square kilometers) in the transition zone of the southeastern United States. Most tall fescue in the US is infected with a symbiotic fungus, *Epichloë coenophiala*, which confers biotic and abiotic stress tolerance for the plant but also produces toxic alkaloids that harm livestock. Although there has been prior evidence that the grass host can influence alkaloid production levels, these have never been precisely quantified. Here, we report on testing alkaloid production and relative fungal biomass on >1000 genetically distinct tall fescue plants. We find that these two traits are weakly correlated, and that both show evidence of moderate to high influence by the host genome. Genome-wide association find only a single marginally significant hit, however, implying that any genetic control by the host is spread among a large number of genes. These results indicate that the host plant exerts moderate influence on these endophytic traits, that the two are largely independent of each other, and that the host’s influence is likely due to a large number of genes of small effect. These results have relevance for breeding tall fescue for forage and turf production, and especially for optimizing the endophyte relationship for tall fescue management.

## 1. Introduction

Tall Fescue (*Lolium arundinaceum*) is an important forage and turf grass that covers 35 million acres (140,000 square kilometers) in the transition zone of the southeastern United States (Hmielowski, 2016). Tall fescue is valued because of its high heat tolerance, adaptability to many soil types, high forage production, and ability to survive heavy grazing from cattle (Breuillin-Sessoms & Watkins, 2020; Jiang & Huang, 2001; Morris, 2010; Novello et al., 2025; Richard et al., 2020).

The most widely planted cultivar of tall fescue in the United States is Kentucky 31, which was released in 1942, and is valued for its widespread adaptation and hardiness (Fergus & Buckner, 1972; Hmielowski, 2016). One of the reasons that Kentucky 31 is so hardy is that it contains a fungal endophyte, *Epichloë coenophiala. E. coenophiala* is an asexually reproducing fungus that propagates by growing within its tall fescue host and infecting developing seeds ensuring it is vertical transmission to the next generation (Christensen et al., 2008; Clay & Schardl, 2002; Zhang et al., 2017).

### 1.1 Effect of *Epichloë coenophiala* on Tall fescue

Fungi in the genus *Epichloë* are thought to help grasses grow in times of drought and environmental stress (Bacon, 1993; Islam et al., 2022; McCulley et al., 2014; Xu et al., 2017). *E. coenophiala* defends its host from herbivores such as insects, nematodes, and mammals (Bacetty et al., 2009; Clay, 1991; Liebe & White, 2018) due to producing various alkaloids (Schardl et al., 2006) that are only produced when inside a plant host (Young et al., 2005). Endophyte free tall fescue lacks persistence in the field and shows a clear decrease in stress tolerance when compared to its endophyte infected counterpart. (J. H. Bouton et al., 1993).

*E. coenophiala* produces several classes of alkaloids, including indole-diterpenes, pyrrolopyrazines (e.g. peramine) , pyrrolizidines (e.g. N-formyllolines), and ergot alkaloids (Takach & Young, 2014). Peramine and the lolines are most well known for their anti-insect properties (Nelli & Scheerer, 2016; Niu et al., 2023; Schardl et al., 2007). Indole-diterpenes best known for their mammalian mycotoxins but can also be insecticidal. However, of these alkaloids, the most relevant for detrimental effects on grazing mammals is ergot alkaloids, which cause vasoconstriction within cattle, which can lead to the loss of weight, gangrenous lesions in extremities, spontaneous abortion, and other aspects of “fescue toxicosis” (Gunter & Beck, 2004). Ergot alkaloid production is encoded by the *EAS* gene cluster (Fleetwood et al., 2007; Florea et al., 2017), and isolates that lack key genes in this cluster have been identified and introduced into tall fescue to produce “non-toxic” varieties (J. Bouton et al., 2002; Hopkins et al., 2010).These “novel” non-toxic isolates do not produce ergot alkaloids. However, due to the high cost of pasture replacement, decades of toxic fescue use, and the added expense of novel endophyte seed, much of the tall fescue in the U.S. remains the toxic variety (Ball et al., 2019; Hmielowski, 2016).

### 1.2 Factors affecting fungal growth and alkaloid production

Studying the mechanisms that control alkaloid production has been difficult. In part this is because the *Epichloë* gene clusters that encode for alkaloids can create a wide range of alkaloid diversity profiles, even within the same species (Bony et al., 2001; Young et al., 2015). *E. coenophiala* can also only reliably produce alkaloids when living inside a host (Young et al., 2005). Nonetheless, some factors have been identified that appear to influence alkaloids *in plantae*. For example, alkaloid levels increase when nitrogen fertilizer is applied, when seed heads have sprouted, during the months of April-May, and in August-September (Hemken et al., 1981; Lea & Smith, 2021; McCulley et al., 2014; Repussard et al., 2014). There is some evidence that the plant influences the amount of alkaloids that *Epichloë* produces (Agee & Hill, 1994; Easton et al., 2002; Faville et al., 2015; West, 2007).

### 1.2 Experimental Aims

We report here the analysis of the interaction between tall fescue and its fungal endophyte, *Epichloë coenophiala*. Our specific goal was to determine the degree to which the plant host can affect fungal growth and alkaloid production *in plantae*, using a pair of biparental mapping populations sharing a common parent and a larger open cross among 17 parents. We report on their estimated heritability correlations between them, and a lack of clear genomic regions in tall fescue influencing these traits. With 1083 genetically distinct individuals tested for both alkaloids and fungal biomass, this represents the largest direct comparison of alkaloid production in tall fescue to date.

## 2. Methods

### 2.1 Population Design

The materials for this study were created by an open cross among 17 tall fescue genotypes, all of which were infected with a similar strain of *Epichloë coenophiala* known to produce ergot alkaloids, peramine and lolines. Production of ergovaline was confirmed in all 17 genotypes with ELISA, SKU number ENDO899-2t (Hill & Agee, 1994). Once the 17 genotypes were established, they were allowed to open pollinate among themselves. Seeds were collected from each plant in 2015, 2016, and 2017, and in 2019 these seeds were planted in a greenhouse at the University of Georgia. Approximately 176 plants per maternal parent were planted, and all plants were tested for endophyte with Agronostics immunoblot kit (Hiatt III et al., 1999); plants without endophyte were discarded.

Initial investigations focused on the entire open cross progeny. Later investigations focused on just the two largest crosses, which formed two half-sib families within this population, totaling 175 progeny and 3 parents.

### 2.2 Sampling

To get the plant genotypes, we extracted DNA from the grass by sampling three leaf blades and freeze-drying them. For the ergot alkaloid measurements, 9 whole tillers were sampled, and leaf blades were discarded. For relative biomass ratios, the grass was sampled from the bottom 1.5 – 2 inches of 3 to 4 tillers where the fungus is most abundant (Figure 1). Both the ergot alkaloid samples and the relative biomass samples were taken once in February of 2023 for the entire population and again in February 2024 for only the two half-sib families.

**Figure 1.**
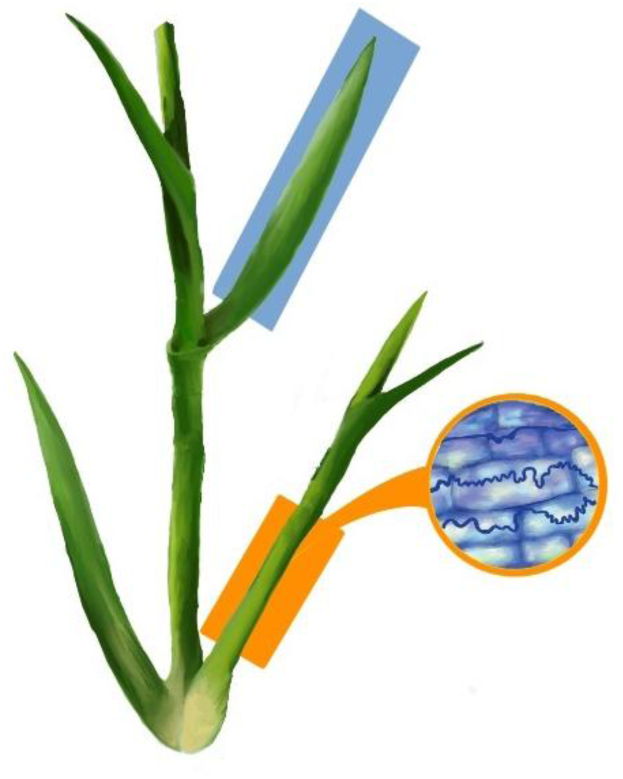
Tall Fescue sampling. Leaf blades (blue box) were sampled for genotyping, while pseudostem (orange box) was taken for alkaloid analysis and fungal biomass since the fungus is concentrated here (Christensen et al., 2008). Figure courtesy of Chloe Mootz.

### 2.3 Fungal Preliminary Data

Differences among original parental lines were tested by selecting 7 parent plants (301, 305, 312, 314, 315, 319, 320) from the 17 founders and creating 5 subclones for each of them. Clones were grown in the greenhouse in a randomized design, each clone was sampled once for both alkaloid testing and the biomass testing, with one random plant sampled twice to evaluate technical variation. Parents 315 and 320 had plants die, so an additional technical replicate was taken from another clone. ANOVA was used to determine if any differences existed among the genotypes, followed by Tukey’s Honest Significant Difference test to determine the differences among them. Similar ANOVA and HSD tests were used later to determine differences among the two half-sib families and the entire population.

### 2.4 Sequencing and Genotyping

Since a reference genome for tall fescue was not available at the time, we performed skim sequencing on the 17 parental lines at a depth of 5x using the Illumina platform (BioProject: PRJNA648970). These sequences were used to create a rough assembly for later mapping. Specifically, the reads were trimmed using TrimGalore (Krueger et al., 2021), and then using Kraken2 (Wood et al., 2019), any reads flagged as non-plant species were removed. De-novo assemblies of the parental lines were created using SPAdes (Bankevich et al., 2012). From these parental assemblies, we chose the most complete one to use as a reference for the next steps. Using BWA (Li & Durbin, 2009), BCF tools (Li, 2011), and VCF tools (Danecek et al., 2011) a VCF file was created and filtered for SNPs with a minor allele frequency of 5%, a mean depth between 20-1000, a minimum quality score of 50, and had to appear in 95% of the parents. The full commands for each of these steps is available at https://github.com/DarrianTalamantes/Genome_creation.git.

From the parental SNPs identified this way, a FlexSeq panel (Rapid Genomics, Gainsville, FL) was developed for use on the progeny and 3000 progeny were tested with this panel.

Because only the maternal parent was known for each progeny, paternal parents were identified based off of shared k-mers. K-mers were tabulated with KMC3 (Kokot et al., 2017) and filtered for ones that were unique to a single parent and appeared in at least six progeny. From this reduced list, the number of unique k-mers that were shared between each possible parent-progeny pair were calculated. Then, for each progeny, the counts of shared k-mers with each possible parent converted into z-scores based on the distribution across all possible parents so that they could be compared across progeny despite different read depths. Z-scores were then clustered using k-means clustering (k=4 clusters) and resampled 100 times. Parentage was assigned when (1) the potential parent-progeny pair appeared in the top cluster (with the highest number of shared SNPs/k-mers) at least 65 out of 100 times; (2) exactly 2 parents passed criterion #1; and (3) one of those two parents was the known maternal parent. Using these cutoffs, we were able to infer parentage for 2407 of the 3000 genotyped progeny.

Later, the two largest crosses, 314x310 and 314x312, and all 17 parents were re-genotyped to greater depth using tunable genotype by sequencing (tGBS)(Ott et al., 2017) outsourced to Freedom Markers. SNPs were called using a draft scaffold-level Tall Fescue genome (Bushman et al. under preparation).

tGBS SNPs were called by quality trimming the reads and aligning them the tall fescue draft genome using GSNAP (Wu & Nacu, 2010); only reads that aligned uniquely were used to identify polymorphic markers. This resulted in a VCF file with 896,240 SNPs. VCFtools (Danecek et al., 2011) and BCFtools (Danecek et al., 2011) were then used to filter SNPs to a minimum allele frequency of 0.05, minimum site depth of 8, maximum site depth of 30, and maximum 50% missing across samples. This resulted in a final genotype set of 12,103 SNPs with an average read depth of 15.9, compared to the average depth of 4.3 for the original Flex-Seq data.

To find genetic outliers, we conducted a PCA on the filtered SNP dataset. Most individuals from each cross grouped with members of their own cross; however, some individuals from 314x312 separated from the main grouping, likely due to misassignment of parentage. Those individuals were filtered out, as well as any individuals that were separate from the main population cluster.

### 2.5 Phenotyping

Total ergot alkaloids were measured via quantitative ELISA as a fee-for-service via the company Agronostics (Schnitzius et al., 2001). Fungal biomass was estimated based on relative DNA amounts for the fungus and tall fescue hosts, measured by quantitative polymerase chain reaction (qPCR). Tillers were harvested as described in section 2.2 and DNA extracted with ZYMO fungal/bacterial miniprep kit (Zymo #D3024) according to the manufacturer’s instructions. Extracted DNA was diluted to 10 ng/µl.

All qPCR reactions used 10 µl of SYBR Green master mix (ROCHE # 04707516001), 1 µl of forward and reverse primers at 10 nm, 3 µl of nuclease free water, and 5 µl of the diluted sample. Using primer sets G3P4 targeting tall fescue glyceraldehyde-3-phosphate dehydrogenase and DMAW4 targeting *Epichloë* dimethylallyl-tryptophan synthase (Talamantes et al., 2024), all reactions were run in triplicate on a Roche 480 II machine with the following settings: pre-incubation at 95°C for 5 minutes, followed by 45 cycles of 95°C for 10 seconds, 50°C for 15 seconds, and 72°C for 16 seconds. The CT values were measured for *Epichloë* and tall fescue to use as a proxy for the biomass of each organism. Any CT values that were 3 standard deviations from the mean for their respective qPCR plate or were measured as 0 were removed before averaging.

Every run had a standard curve made of tenfold serial dilutions (5 concentrations) of purified PCR product. G3P4 product initial concentration was 0.001 ng/µl, and DMAW4 was 0.01 ng/µl. Each run had two sets of serially diluted product in case one standard curve failed. These standard curves allowed for the calculation of qPCR efficiency and r^2^ of the standard curve. Any runs with standard curves with r^2^ values below 0.95 or primer efficiency below 85% were redone. The averaged CT values were normalized and adjusted for primer amplification efficiency according to the following equation:

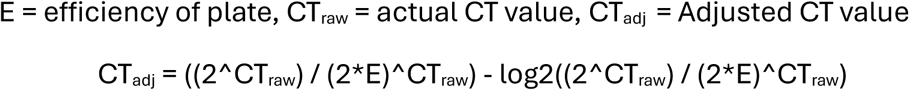

To get the efficiency-adjusted CT ratio, the efficiency-adjusted CT value for *Epichloë* was divided by the efficiency-adjusted CT value for tall fescue.

Phenotyping was done for 1083 of the 3000 total individuals. We did not phenotype individuals that we could not identify parentage or that had died between the genotyping in 2019 and phenotyping in 2023. Phenotyping was done one more time for the two half-sib families in 2024. After filtering, the two half-sib families totaled 133 individuals.

Residuals for the data were then taken by running the lm() function in R and removing known environmental factors (sampling date for alkaloids, and sampling date and assay plate for qPCR). For the half-sib families, the phenotypes were averaged across the two years they were taken, and then the residuals were found as above.

All scripts are available at https://github.com/DarrianTalamantes/Mapping_and_QTL/tree/main/Non_snakemake_workflow/R_Files.

### 2.6 Heritability Calculation and GWAS

Heritability was estimated for the full population using the Flex Seq genotypes and the phenotypes taken in 2023. For the half-sib population, Heritability was calculated using the Freedom Markers genotypes, and a separate calculation was made for each set of phenotypes: 2023, 2024, and the average of both years. First, TASSEL (Bradbury et al., 2007) was used to convert the filtered VCF file into a numerical genotype file. The numerical genotype file was then used in rrBLUP (Endelman, 2011) to create an additive kinship matrix. The kinship matrix and the phenotypes were then used to do genotypic value prediction based on kinship. The equation below was used to calculate heritability.

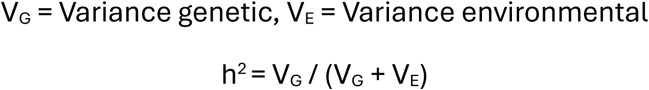

GWAS was conducted on the half-sib families with TASSEL. First, both principal components and a kinship matrix were created from the filtered SNPs (section 2.4). The genotypes, first 5 PCs, and residual phenotype files were joined and combined with the kinship matrix to run a mixed linear model (MLM). The MLM was also run with the separated 2023 and 2024 residual phenotypes. P-value cut-offs for GWAS were calculated using both the false discovery rate and the Bonferroni correction.

A GWAS and heritability calculations were also run on the larger population in a similar manner, using the 1083 individuals that we had phenotypes and genotypes for.

## 3. Results

### 3.1 Phenotypic relationship between alkaloids and fungal biomass

Seven of the 17 parental lines were selected to determine how fungal biomass and alkaloid concentrations varied by genotype. There were significant differences among these lines for total ergot alkaloids but not for relative fungal biomass (Figure 2). Phenotypes among the parental lines were weakly but significantly correlated (r^2^= 0.132; p = 0.02).

**Figure 2:**
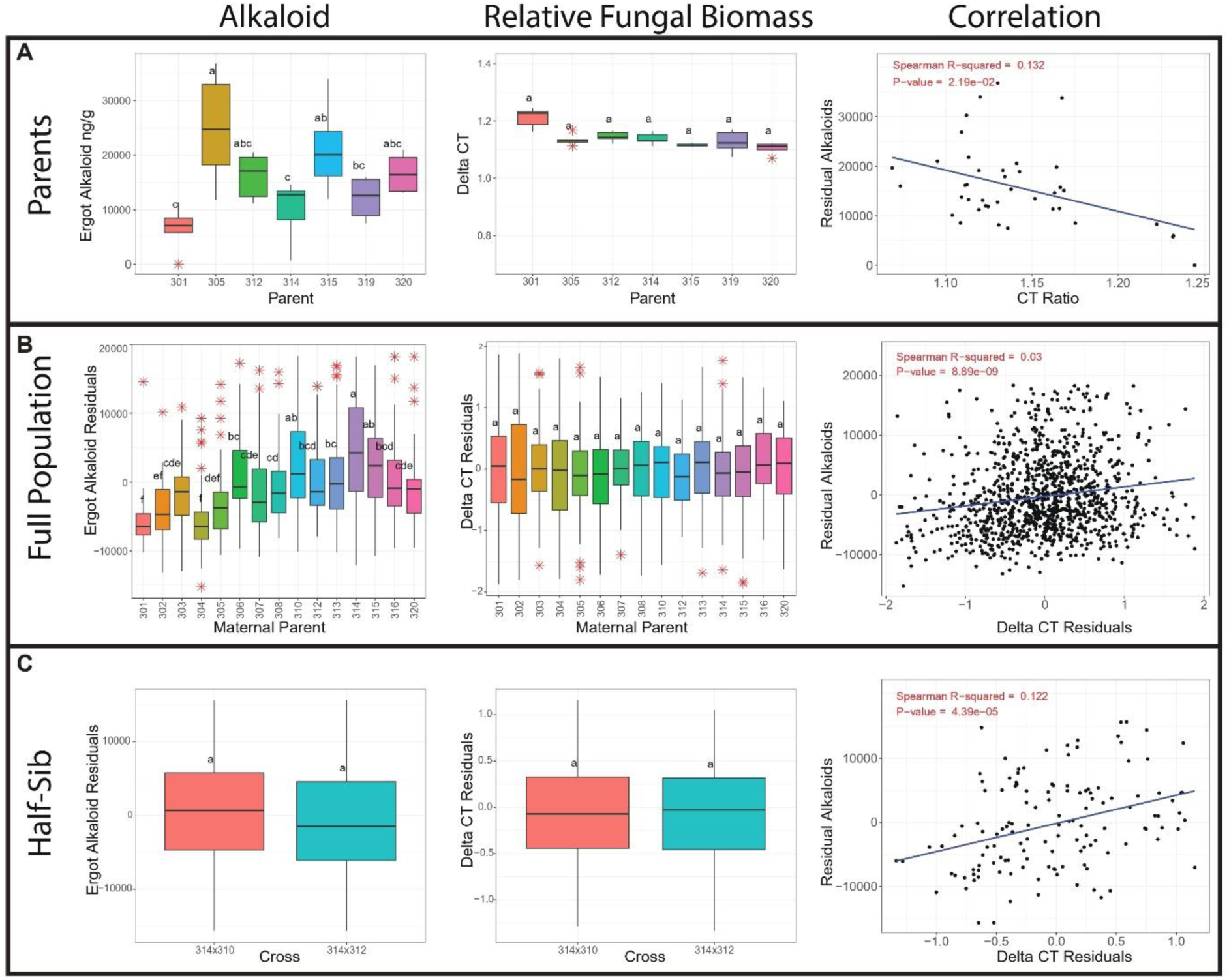
The relationship between alkaloid production and fungal biomass. Analyses are shown for a subset of parents (A), the entire population (B), and the two half-sib families for which additional data was collected (C). (Boxplots in B are separated based on maternal parent since the endophyte is only inherited maternally.) For each population group, the leftmost plot shows the amount of ergot alkaloids present as measured by ELISA. (Actual values in A, and residual values in B & C after fitting a statistical model to correct for batch effects.) The center column shows relative fungal biomass, as measured by the difference in crossing threshold (“delta CT”) of qPCR for either the plant or the fungus. (B and C again use residual instead of raw values to correct for batch effects.) The correlations between alkaloids and fungal biomass are shown at right.

In 2023, 1083 individuals from the overall mapping population and the 17 parents were phenotyped for total ergot alkaloid concentration and relative fungal biomass. To estimate fungal biomass, a delta CT ratio of fungus/plant PCR product was used. This data was filtered for outliers and normalized (see Methods), then grouped by maternal parent. This approach was chosen because the maternal parentage is known, while paternal parentage was inferred. Additionally, grouping by maternal lineage ensures consistency in *Epichloë* presence across groups, as the endophyte is maternally inherited. We find significant differences in alkaloid production among the maternal groups but no differences in the relative fungal biomass data (Figure 2). The correlation between these phenotypes is numerically small (r^2^ = .03) but highly significant (p=8.89 x 10^-9^), likely due to the large sample size.

As a focused test, we selected out the two largest crosses (based on known maternal and genotype-inferred paternal parents) for deeper genotyping and a second round of phenotyping. These families share a common parent (314) and so are half-sib families. We did not detect any significant differences in phenotype for these families (Figure 2), which again show a statistically significant but numerically small relationship between fungal biomass and alkaloid production (r^2^=0.122, p=4.39 x 10^-5^) (Figure 2). This pattern holds when individual timepoints are analyzed separately (Figure 3). When comparing across timepoints, correlations for both traits were relatively low (r^2^= 0.063; p = 3.06x10^-3^ for alkaloids and r^2^= 0.133; p = 9.78x10^-6^ for relative biomass).

**Figure 3:**
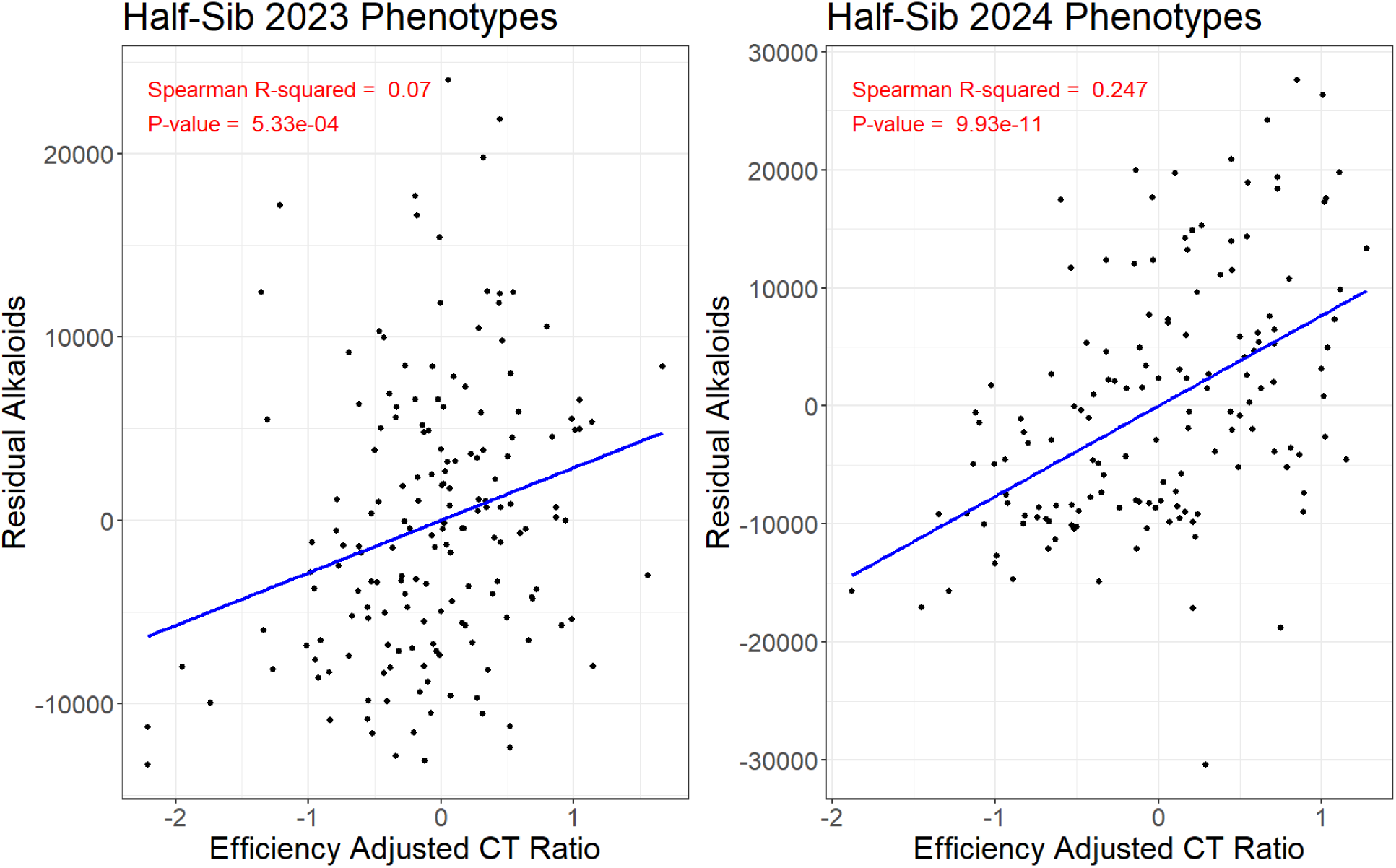
Correlation between the half-sib population phenotypes. Determination of alkaloid and CT ratios is the same as in Figure 2C, save that the data for the two sampling years (2023 and 2024) were kept separate. Both years show statistically significant correlations between fungal biomass and alkaloid production, though the relationship is much stronger in 2023.

### 3.2 Heritability estimates of alkaloid production and fungal biomass

“Heritability” refers to the amount of variability in a trait that is explainable by genetics (specifically the genetics of the host plant in this study). Narrow-sense heritability for the overall population was calculated using genetic and trait data for 1083 individuals, finding an estimated h^2^ of 0.352 for ergot alkaloids and 0.672 for relative fungal biomass.

Genome-wide association was conducted on these individuals, which resulted in a single significant SNP (FaChr7G1: 214311125) for relative biomass (Figure 4). The 25kb segment up- and downstream of this SNP were BLASTed against the *Lolium Perenne* (Frei et al., 2021) and rice (Shang et al., 2023) genomes to identify potential candidate genes, but only uncharacterized proteins were found (Table S1).

**Figure 4:**
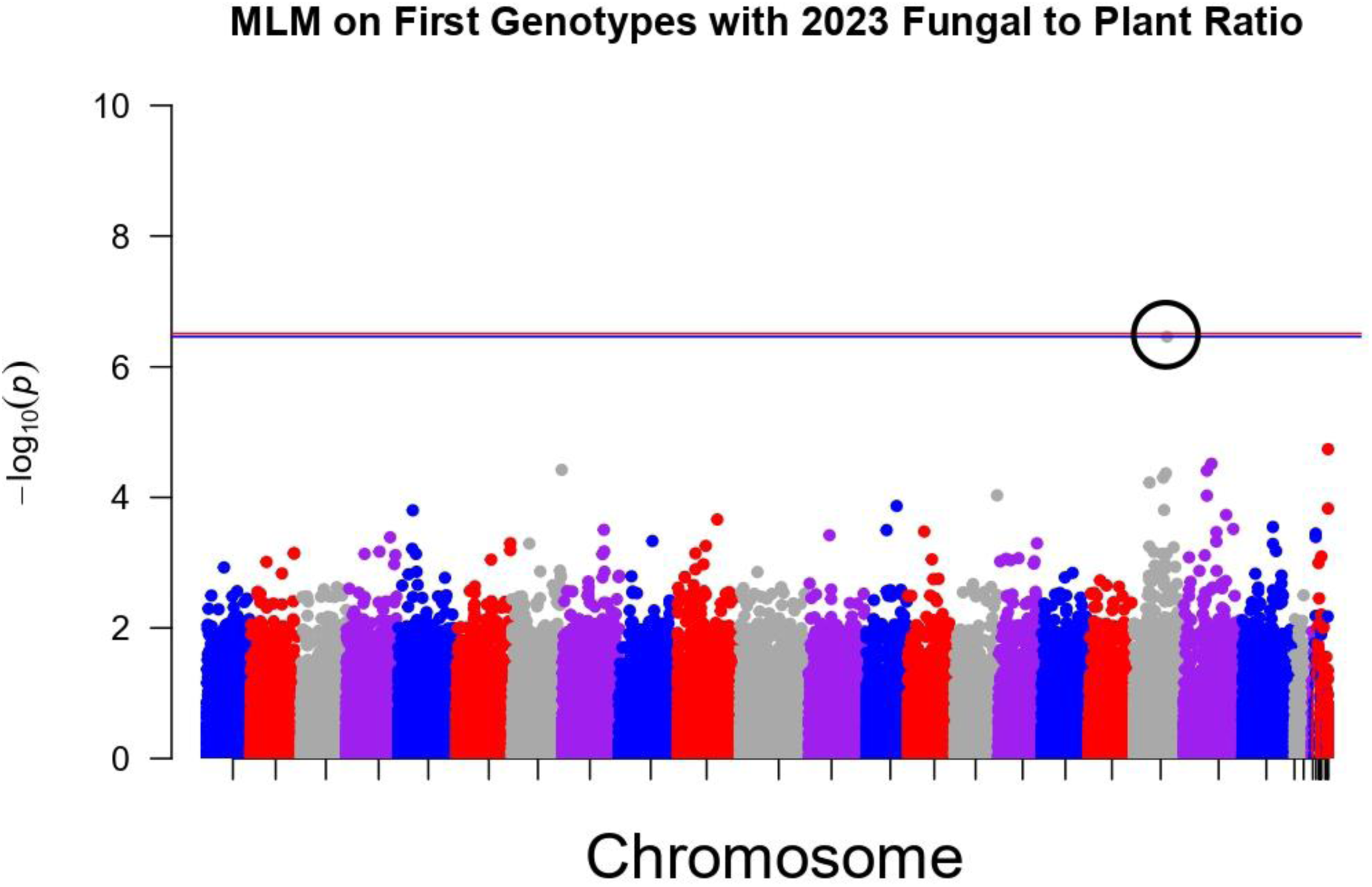
Manhattan plot of genome-wide association for relative fungal biomass. A mixed linear model was fit in TASSEL (Bradbury et al. 2007) for relative biomass (delta CT) across 1083 individuals. Only a single SNP (circled, at FaChr7G1: 214311125) passes the 5% False Discovery Rate cutoff (blue), though not the Bonferroni-corrected cutoff (red).

Narrow sense heritability was similarly estimated for the values of the half-sib families. When calculated for both years, heritability was 0.603 for alkaloid concentration and 0.702 for relative fungal biomass. In 2023, heritability was lowest for both traits (0.245 for alkaloids and 0.171 for relative fungal biomass), while in 2024, heritability increased to 0.465 for alkaloids and 0.730 for relative fungal biomass. Genetic mapping of these data sets found no sites that passed a 5% false discovery rate threshold (data not shown).

## 4. Discussion and Conclusions

In this study, we compared the relative fungal biomass and alkaloid production of over 1000 genetically unique tall fescue plants grown in a common greenhouse environment. Our principal aims were to quantify the relationship between these traits and the degree to which both are influenced by the genome of their host. Our results support the following conclusions:

### 4.1 Fungal Biomass is Weakly Correlated to Alkaloid Production

Across all population groups, the relative biomass of *Epichloë* was weakly but significantly correlated with fungal alkaloid production. We found that in the half-sib families, the correlation varied ∼3.5x in two consecutive years (2023 and 2024; 0.07 vs 0.25), implying that environmental or other factors may change the relationship between the traits. Alkaloid levels tend to respond to environmental variability (Dinkins et al., 2023); if fungal biomass is more constant, then it could explain the variable relationship between these traits. Our data indicate that fungal biomass is marginally more stable (r^2^ of 0.133 vs 0.063), though the correlation is still weak. This low stability of fungal biomass may be due to the host continually producing new tillers, which would allow for different rates of fungal infection over time.

Other studies that have investigated the correlation between *Epichloë* biomass and alkaloid concentrations consistently have a higher R^2^ values (Cagnano et al., 2020; Fuchs et al., 2017b), though they are also looking at related species (*Lolium perenne* and *Epichloë festucae*) and not tall fescue/*Epichloë coenophiala* specifically. Differences may also be due to methodology. Liquid chromatography/mass spectrometry on >1000 individuals proved to be logistically unfeasible, hence why we chose ELISA quantification and qPCR (Talamantes et al., 2024), the latter of which is similar to how *Epichloë* festucae was measured within *Lolium perenne*. (Fuchs et al., 2017a).

### 4.2 Both Biomass and Alkaloid Production Show Evidence of Plant Genetic Influence

Although the two traits examined here (alkaloid production and fungal biomass) show only weak relationship to each other, they both show a moderate relationship to the genetics of their host plant. Evidence of host genetic control has been shown before (Agee & Hill, 1994), and our results are the largest and most thorough quantification of this effect to date. Both ergot alkaloid concentration and relative fungal biomass were found to be moderately heritable traits within tall fescue, with values ranging from 0.245-0.603 for alkaloid concentration and from 0.171-0.730 for relative fungal biomass. With one exception (half-sib families in 2023, with the lowest heritability overall), fungal biomass was consistently more heritable than alkaloid production.

Previous studies that have measured heritability have relied on phenotypic data alone and also found alkaloid production and fungal biomass to be relatively heritable traits (e.g., heritabilities of 0.49–0.72) (Adcock et al., 1997; Easton et al., 2002). Our study found similar heritability estimates for both traits, supporting the idea that tall fescue exerts control over fungal phenotypes. Combined with the low R² between traits, this suggests that alkaloid levels and fungal biomass are influenced by plant genetics but likely governed by different genes or genetic networks.

Investigation into possible genes that governed these traits found only one possible SNP, at FaChr7G1: 214311125. However, no protein encoding genes with known functions were identified within 25kb of this SNP. Combined with the moderate heritability results of 0.35 (alkaloids) and 0.67 (fungal biomass), these results imply that the host genetic influence on these traits is highly quantitative and diffuse, meaning that many genes across the genome are each likely contributing a small effect. These results contrast with a QTL analysis on the same traits of alkaloid amounts and fungal biomass with perennial ryegrass and *Epichloë festucae*, which yielded 11 genomic regions affecting these traits (Faville et al., 2015). Given the close relationship between tall fescue and perennial ryegrass, it is curious that we find such different results; one possibility is that since perennial ryegrass is a diploid species whereas tall fescue is hexaploid, the latter’s genetic effects are simply spread among its 3 genome copies and thus more difficult to identify.

### 4.3 Outlook and Future Directions

Tall fescue covers 140,000 km^2^ of the southern US, and ergot alkaloid toxicity is estimated to cost US growers between $600 million to $1 billion per year (Hancock & Andrae, 2017). Although newer cultivars now exist that eliminate livestock toxicity, adoption has been slow. (J. Bouton et al., 2002). Most pastures still consist primarily of toxic cultivars because remediation is costly and non-toxic cultivars require more management, as cattle tend to overgraze them (Hmielowski, 2016); efforts to accelerate this transition through public-private partnerships and education are underway and showing significant promise (Roberts et al., 2020) Understanding the mechanisms that control both fungal presence and alkaloid production will provide insight into this intimate and highly beneficial symbiosis, but also potentially identify ways to more economically remediate these pastures and/or mitigate the amount of alkaloids produced and ingested by cattle. Ultimately, these results aim to improve the lives of both livestock and ranchers dependent on tall fescue and provide insight for superior management of pasture resources.

## Acknowledgements

This work was supported by NSF grant # 1764127. The authors would like to thank members of the Wallace lab for assistance with sampling fescue plants, especially Naomi Rodman, and Michael Trammell (Oklahoma State University) for the production of tall fescue seeds at the Noble Research Institute, Chloe Mootz for providing Figure 1, and Drs. Shaun Bushman and Matthew Robbins for early access to the tall fescue genome assembly.

## Author Contributions

DRT: Methodology, Investigation, Data Curation, Formal Analysis, Visualization. Writing-Original Draft. CY: Resources, Writing – Review and Editing. JGW: Conceptualization, Writing – Review and Editing, Supervision, Project Administration, Funding acquisition.

---

**Supplementary Table 1:** The top 15 hits of proteins found within 25,000 base pairs from the significant SNP for relative fungal biomass.

| Rice |  |  |  |  |  |  |
| --- | --- | --- | --- | --- | --- | --- |
| Query | Subject (Rice) | Rice protein | % identity | length | e-value | bitscore |
| FaChr7G1:214286125-214336125 | XP_066161687.1 | XP_066161687.1 uncharacterized protein [Oryza sativa Japonica Group] | 63.562 | 1106 | 0 | 1400 |
| FaChr7G1:214286125-214336125 | XP_066168244.1 | XP_066168244.1 uncharacterized protein [Oryza sativa Japonica Group] | 63.472 | 1106 | 0 | 1399 |
| FaChr7G1:214286125-214336125 | XP_066167998.1 | XP_066167998.1 uncharacterized protein [Oryza sativa Japonica Group] | 63.472 | 1106 | 0 | 1399 |
| FaChr7G1:2142861<br>25-214336125 | XP_06616440<br>1.1 | XP_066164401.1<br>uncharacterized<br>protein [Oryza<br>sativa Japonica<br>Group] | 63.47<br>2 | 1106 | 0 | 1399 |
| FaChr7G1:2142861<br>25-214336125 | XP_06616228<br>8.1 | XP_066162288.1<br>uncharacterized<br>protein [Oryza<br>sativa Japonica<br>Group] | 63.47<br>2 | 1106 | 0 | 1399 |
| FaChr7G1:2142861<br>25-214336125 | XP_06616733<br>4.1 | XP_066167334.1<br>uncharacterized<br>protein [Oryza<br>sativa Japonica<br>Group] | 63.38<br>2 | 1106 | 0 | 1395 |
| FaChr7G1:2142861<br>25-214336125 | XP_06616227<br>5.1 | XP_066162275.1<br>uncharacterized<br>protein [Oryza<br>sativa Japonica<br>Group] | 63.38<br>2 | 1106 | 0 | 1394 |
| FaChr7G1:2142861<br>25-214336125 | XP_06616157<br>2.1 | XP_066161572.1<br>uncharacterized<br>protein [Oryza<br>sativa Japonica<br>Group] | 63.38<br>2 | 1106 | 0 | 1393 |
| FaChr7G1:2142861<br>25-214336125 | XP_06616402<br>3.1 | XP_066164023.1<br>uncharacterized<br>protein [Oryza<br>sativa Japonica<br>Group] | 63.37<br>3 | 1103 | 0 | 1390 |
| FaChr7G1:2142861<br>25-214336125 | XP_06616649<br>0.1 | XP_066166490.1<br>uncharacterized<br>protein [Oryza<br>sativa Japonica<br>Group] | 61.23<br>2 | 1104 | 0 | 1334 |
| FaChr7G1:2142861<br>25-214336125 | XP_06616335<br>9.1 | XP_066163359.1<br>uncharacterized<br>protein [Oryza<br>sativa Japonica<br>Group] | 60.92<br>5 | 1103 | 0 | 1318 |
| FaChr7G1:2142861<br>25-214336125 | XP_06616305<br>5.1 | XP_066163055.1<br>uncharacterized<br>protein, partial | 57.29<br>8 | 1103 | 0 | 1206 |
|  |  | [Oryza sativa Japonica Group] |  |  |  |  |
| FaChr7G1:214286125-214336125 | XP_066165204.1 | XP_066165204.1 uncharacterized protein, partial [Oryza sativa Japonica Group] | 57.246 | 1104 | 0 | 1205 |
| FaChr7G1:214286125-214336125 | XP_066166206.1 | XP_066166206.1 uncharacterized protein [Oryza sativa Japonica Group] | 57.156 | 1104 | 0 | 1202 |
| FaChr7G1:214286125-214336125 | XP_066162661.1 | XP_066162661.1 uncharacterized protein [Oryza sativa Japonica Group] | 57.156 | 1104 | 0 | 1198 |
| Lolium Perenne |  |  |  |  |  |  |
| Query | Subject (Rice) | Rice protein | % identity | length | e-value | bitscore |
| FaChr7G1:214286125-214336125 | XP_071677341.1 | XP_071677341.1 uncharacterized protein [Lolium perenne] | 94.316 | 1038 | 0 | 2034 |
| FaChr7G1:214286125-214336125 | XP_051185024.1 | XP_051185024.1 uncharacterized protein [Lolium perenne] | 73.622 | 1107 | 0 | 1584 |
| FaChr7G1:214286125-214336125 | XP_071685507.1 | XP_071685507.1 uncharacterized mitochondrial protein AtMg00860-like [Lolium perenne] | 47.465 | 217 | 2.60e-65 | 224 |
| FaChr7G1:214286125-214336125 | XP_071685506.1 | XP_071685506.1 uncharacterized protein [Lolium perenne] | 34.844 | 706 | 6.31e-121 | 424 |
| FaChr7G1:214286125-214336125 | XP_071685506.1 | XP_071685506.1 uncharacterized protein [Lolium perenne] | 40.625 | 96 | 8.38e-09 | 65.1 |
| FaChr7G1:2142861<br>25-214336125 | XP_07168548<br>6.1 | XP_071685486.1<br>uncharacterized<br>protein [Lolium<br>perenne] | 68.65<br>3 | 386 | 7.10<br>e-170 | 559 |
| FaChr7G1:2142861<br>25-214336125 | XP_07168548<br>6.1 | XP_071685486.1<br>uncharacterized<br>protein [Lolium<br>perenne] | 40.09<br>2 | 651 | 3.09<br>e-142 | 478 |
| FaChr7G1:2142861<br>25-214336125 | XP_07168546<br>9.1 | XP_071685469.1<br>uncharacterized<br>protein [Lolium<br>perenne] | 54.31<br>5 | 197 | 9.62<br>e-61 | 214 |
| FaChr7G1:2142861<br>25-214336125 | XP_07168546<br>5.1 | XP_071685465.1<br>uncharacterized<br>protein [Lolium<br>perenne] | 60.10<br>4 | 193 | 1.43<br>e-73 | 248 |
| FaChr7G1:2142861<br>25-214336125 | XP_07168546<br>4.1 | XP_071685464.1<br>uncharacterized<br>protein [Lolium<br>perenne] | 71.42<br>9 | 210 | 1.06<br>e-93 | 308 |
| FaChr7G1:2142861<br>25-214336125 | XP_07168546<br>1.1 | XP_071685461.1<br>uncharacterized<br>protein [Lolium<br>perenne] | 37.89<br>8 | 628 | 3.56<br>e-134 | 442 |
| FaChr7G1:2142861<br>25-214336125 | XP_07168546<br>0.1 | XP_071685460.1<br>uncharacterized<br>protein [Lolium<br>perenne] | 74.92<br>4 | 654 | 0 | 1053 |
| FaChr7G1:2142861<br>25-214336125 | XP_07168546<br>0.1 | XP_071685460.1<br>uncharacterized<br>protein [Lolium<br>perenne] | 60.80<br>4 | 199 | 3.80<br>e-65 | 249 |
| FaChr7G1:2142861<br>25-214336125 | XP_07168546<br>0.1 | XP_071685460.1<br>uncharacterized<br>protein [Lolium<br>perenne] | 36.50<br>8 | 126 | 1.07<br>e-07 | 61.2 |
| FaChr7G1:2142861<br>25-214336125 | XP_07168545<br>3.1 | XP_071685453.1<br>uncharacterized<br>protein [Lolium<br>perenne] | 36.84<br>2 | 171 | 4.33<br>e-27 | 112 |

## References

1. Adcock, R., Hill, N., Bouton, J., Boerma, H., & Ware, G. (1997). Symbiont Regulation and Reducing Ergot Alkaloid Concentration by Breeding Endophyte-Infected Tall Fescue. Journal of Chemical Ecology - J CHEM ECOL, 23, 691–704. 10.1023/B:JOEC.0000006404.33191.60

2. Agee, C., & Hill, N. (1994). Ergovaline Variability in Acremonium-Infected Tall Fescue Due to Environment and Plant Genotype. Crop Science - CROP SCI, 34. 10.2135/cropsci1994.0011183X003400010040x

3. Bacetty, A. A., Snook, M. E., Glenn, A. E., Noe, J. P., Hill, N., Culbreath, A., Timper, P., Nagabhyru, P., & Bacon, C. W. (2009). Toxicity of endophyte-infected tall fescue alkaloids and grass metabolites on Pratylenchus scribneri. Phytopathology, 99(12), 1336–1345. 10.1094/PHYTO-99-12-1336

4. Bacon, C. W. (1993). Abiotic stress tolerances (moisture, nutrients) and photosynthesis in endophyte-infected tall fescue. *Agriculture*, Ecosystems & Environment, 44(1), 123–141. 10.1016/0167-8809(93)90042-N

5. Ball, D. M., Lacefield, G. D., & Hoveland, C. S. (2019). The Wonder Grass: The Story of Tall Fescue in the United States. https://aurora.auburn.edu/handle/11200/49449

6. Bankevich, A., Nurk, S., Antipov, D., Gurevich, A. A., Dvorkin, M., Kulikov, A. S., Lesin, V. M., Nikolenko, S. I., Pham, S., Prjibelski, A. D., Pyshkin, A. V., Sirotkin, A. V., Vyahhi, N., Tesler, G., Alekseyev, M. A., & Pevzner, P. A. (2012). SPAdes: A New Genome Assembly Algorithm and Its Applications to Single-Cell Sequencing. Journal of Computational Biology, 19(5), 455–477. 10.1089/cmb.2012.0021

7. Bony, S., Pichon, N., Ravel, C., Durix, A., Balfourier, F., & Guillaumin, J.-J. (2001). The relationship between mycotoxin synthesis and isolate morphology in fungal endophytes of Lolium perenne. New Phytologist, 152(1), 125–137. 10.1046/j.0028-646x.2001.00231.x

8. Bouton, J. H., Gates, R. N., Belesky, D. P., & Owsley, M. (1993). Yield and Persistence of Tall Fescue in the Southeastern Coastal Plain after Removal of Its Endophyte. Agronomy Journal, 85(1), 52–55. 10.2134/agronj1993.00021962008500010011x

9. Bouton, J., Latch, G., Hill, N., Hoveland, C., McCann, M., Watson, R., Parish, J., Hawkins, L., & Thompson, F. (2002). Reinfection of Tall Fescue Cultivars with Non-Ergot Alkaloid– Producing Endophytes. Agronomy Journal - AGRON J, 94. 10.2134/agronj2002.0567

10. Bradbury, P. J., Zhang, Z., Kroon, D. E., Casstevens, T. M., Ramdoss, Y., & Buckler, E. S. (2007). TASSEL: Software for association mapping of complex traits in diverse samples. Bioinformatics, 23(19), 2633–2635. 10.1093/bioinformatics/btm308

11. Breuillin-Sessoms, F., & Watkins, E. (2020). Performance of multiple turfgrass species during prolonged heat stress and recovery in a controlled environment. Crop Science, 60(6), 3344–3361. 10.1002/csc2.20262

12. Cagnano, G., Lenk, I., Roulund, N., Jensen, C. S., Cox, M. P., & and Asp, T. (2020). Mycelial biomass and concentration of loline alkaloids driven by complex population structure in Epichloë uncinata and meadow fescue (Schedonorus pratensis). Mycologia, 112(3), 474–490. 10.1080/00275514.2020.1746607

13. Christensen, M. J., Bennett, R. J., Ansari, H. A., Koga, H., Johnson, R. D., Bryan, G. T., Simpson, W. R., Koolaard, J. P., Nickless, E. M., & Voisey, C. R. (2008). Epichloë endophytes grow by intercalary hyphal extension in elongating grass leaves. Fungal Genetics and Biology, 45(2), 84–93. 10.1016/j.fgb.2007.07.013

14. Clay, K. (1991). Endophytes as Antagonists of Plant Pests. In J. H. Andrews & S. S. Hirano (Eds.), Microbial Ecology of Leaves (pp. 331–357). Springer. 10.1007/978-1-4612-3168-4_17

15. Clay, K., & Schardl, C. (2002). Evolutionary origins and ecological consequences of endophyte symbiosis with grasses. The American Naturalist, 160 *Suppl 4*, S99– S127. 10.1086/342161

16. Danecek, P., Auton, A., Abecasis, G., Albers, C. A., Banks, E., DePristo, M. A., Handsaker, R. E., Lunter, G., Marth, G. T., Sherry, S. T., McVean, G., Durbin, R., & 1000 Genomes Project Analysis Group. (2011). The variant call format and VCFtools. Bioinformatics, 27(15), 2156–2158. 10.1093/bioinformatics/btr330

17. Dinkins, R. D., Coe, B. L., Phillips, T. D., & Ji, H. (2023). Accumulation of Alkaloids in Different Tall Fescue KY31 Clones Harboring the Common Toxic Epichloë coenophiala Endophyte under Field Conditions. Agronomy, 13(2), Article 2. 10.3390/agronomy13020356

18. Easton, H. S., Latch, G. C. M., Tapper, B. A., & Ball, O. J.-P. (2002). Ryegrass Host Genetic Control of Concentrations of Endophyte-Derived Alkaloids. Crop Science, 42(1), 51–57. 10.2135/cropsci2002.5100

19. Endelman, J. B. (2011). Ridge Regression and Other Kernels for Genomic Selection with R Package rrBLUP. The Plant Genome, 4(3). 10.3835/plantgenome2011.08.0024

20. Faville, M. J., Briggs, L., Cao, M., Koulman, A., Jahufer, M. Z. Z., Koolaard, J., & Hume, D. E. (2015). A QTL analysis of host plant effects on fungal endophyte biomass and alkaloid expression in perennial ryegrass. Molecular Breeding: New Strategies in Plant Improvement, 35(8), 161. 10.1007/s11032-015-0350-1

21. Fergus, E. N., & Buckner, R. C. (1972). Registration of Kentucky 31 Tall Fescue (Reg. No. 7). Crop Science, 12(5), cropsci1972.0011183X001200050061x. 10.2135/cropsci1972.0011183X001200050061x

22. Fleetwood, D. J., Scott, B., Lane, G. A., Tanaka, A., & Johnson, R. D. (2007). A Complex Ergovaline Gene Cluster in Epichloë Endophytes of Grasses. Applied and Environmental Microbiology, 73(8), 2571–2579. 10.1128/AEM.00257-07

23. Florea, S., Panaccione, D. G., & Schardl, C. L. (2017). Ergot Alkaloids of the Family Clavicipitaceae. Phytopathology®, 107(5), 504–518. 10.1094/PHYTO-12-16-0435-RVW

24. Frei, D., Veekman, E., Grogg, D., Stoffel-Studer, I., Morishima, A., Shimizu-Inatsugi, R., Yates, S., Shimizu, K. K., Frey, J. E., Studer, B., & Copetti, D. (2021). Ultralong Oxford Nanopore Reads Enable the Development of a Reference-Grade Perennial Ryegrass Genome Assembly. Genome Biology and Evolution, 13(8), evab159. 10.1093/gbe/evab159

25. Fuchs, B., Krischke, M., Mueller, M. J., & Krauss, J. (2017a). Herbivore-specific induction of defence metabolites in a grass–endophyte association. Functional Ecology, 31(2), 318–324. 10.1111/1365-2435.12755

26. Fuchs, B., Krischke, M., Mueller, M. J., & Krauss, J. (2017b). Plant age and seasonal timing determine endophyte growth and alkaloid biosynthesis. Fungal Ecology, 29, 52–58. 10.1016/j.funeco.2017.06.003

27. Gunter, S. A., & Beck, P. A. (2004). Novel endophyte-infected tall fescue for growing beef cattle. Journal of Animal Science, 82(suppl_13), E75–E82. 10.2527/2004.8213_supplE75x

28. Hancock, D. W., & Andrae, J. (2017, March). Novel Endophyte-Infected Tall Fescue. University of Georgia Extension. https://secure.caes.uga.edu/extension/publications/files/pdf/C%20861_4.PDF

29. Hemken, R. W., Boling, J. A., Bull, L. S., Hatton, R. H., Buckner, R. C., & Bush, L. P. (1981). Interaction of Environmental Temperature and Anti-Quality Factors on the Severity of Summer Fescue Toxicosis. Journal of Animal Science, 52(4), 710–714. 10.2527/jas1981.524710x

30. Hiatt III, E. E., Hill, N. S., Bouton, J. H., & Stuedemann, J. A. (1999). Tall Fescue Endophyte Detection: Commercial Immunoblot Test Kit Compared with Microscopic Analysis. Crop Science, 39(3), cropsci1999.0011183X003900030030x. 10.2135/cropsci1999.0011183X003900030030x

31. Hill, N. S., & Agee, C. S. (1994). Detection of Ergoline Alkaloids in Endophyte-Infected Tall Fescue by Immunoassay. Crop Science, 34(2), cropsci1994.0011183X003400020041x. 10.2135/cropsci1994.0011183X003400020041x

32. Hmielowski, T. (2016). The fascinating tale of tall fescue. CSA News, 61(12), 4–9. 10.2134/csa2016-61-12-1

33. Hopkins, A. A., Young, C. A., Panaccione, D. G., Simpson, W. R., Mittal, S., & Bouton, J. H. (2010). Agronomic Performance and Lamb Health among Several Tall Fescue Novel Endophyte Combinations in the South-Central USA. Crop Science, 50(4), 1552– 1561. 10.2135/cropsci2009.08.0473

34. Islam, M. S., Krom, N., Kwon, T., Li, G., & Saha, M. C. (2022). Transcriptome of Endophyte-Positive and Endophyte-Free Tall Fescue Under Field Stresses. Frontiers in Plant Science, 13. 10.3389/fpls.2022.803400

35. Jiang, Y., & Huang, B. (2001). Physiological Responses to Heat Stress Alone or in Combination with Drought: A Comparison between Tall Fescue and Perennial Ryegrass. 10.21273/HORTSCI.36.4.682

36. Kokot, M., Długosz, M., & Deorowicz, S. (2017). KMC 3: Counting and manipulating k-mer statistics. Bioinformatics, 33(17), 2759–2761. 10.1093/bioinformatics/btx304

37. Krueger, F., James, F., Ewels, P., Afyounian, E., & Schuster-Boeckler, B. (2021). *FelixKrueger/TrimGalore: V0.6.7 - DOI via Zenodo* (Version 0.6.7) [Computer software]. Zenodo. 10.5281/zenodo.5127899

38. Lea, K. L. M., & Smith, S. R. (2021). Using On-Farm Monitoring of Ergovaline and Tall Fescue Composition for Horse Pasture Management. Toxins, 13(10), Article 10. 10.3390/toxins13100683

39. Li, H. (2011). A statistical framework for SNP calling, mutation discovery, association mapping and population genetical parameter estimation from sequencing data. Bioinformatics, 27(21), 2987–2993. 10.1093/bioinformatics/btr509

40. Li, H., & Durbin, R. (2009). Fast and accurate short read alignment with Burrows-Wheeler transform. *Bioinformatics (Oxford*, England), 25(14), 1754–1760. 10.1093/bioinformatics/btp324

41. Liebe, D. M., & White, R. R. (2018). Meta-analysis of endophyte-infected tall fescue effects on cattle growth rates. Journal of Animal Science, 96(4), 1350–1361. 10.1093/jas/sky055

42. McCulley, R. L., Bush, L. P., Carlisle, A. E., Ji, H., & Nelson, J. A. (2014). Warming reduces tall fescue abundance but stimulates toxic alkaloid concentrations in transition zone pastures of the U.S. Frontiers in Chemistry, 2, 88. 10.3389/fchem.2014.00088

43. Morris, K. (2010). 2006 *National Tall Fescue Test* (National Turfgrass Evaluation Program) [Progress Report]. USDA-ARS.

44. Nelli, M. R., & Scheerer, J. R. (2016). Synthesis of Peramine, an Anti-insect Defensive Alkaloid Produced by Endophytic Fungi of Cool Season Grasses. Journal of Natural Products, 79(4), 1189–1192. 10.1021/acs.jnatprod.5b01089

45. Niu, J., Qi, J., Wang, P., Liu, C., & Gao, J. (2023). The chemical structures and biological activities of indole diterpenoids. Natural Products and Bioprospecting, 13(1), 3. 10.1007/s13659-022-00368-7

46. Novello, A., Pornaro, C., Fidanza, M., & Macolino, S. (2025). Adaptability and Character Traits of Turf-type Tall Fescue Cultivars Grown under Limited Irrigation in Northern Italy. 10.21273/HORTTECH05512-24

47. Ott, A., Liu, S., Schnable, J. C., Yeh, C.-T. ‘Eddy,’ Wang, K.-S., & Schnable, P. S. (2017). tGBS® genotyping-by-sequencing enables reliable genotyping of heterozygous loci. Nucleic Acids Research, 45(21), e178. 10.1093/nar/gkx853

48. Repussard, C., Zbib, N., Tardieu, D., & Guerre, P. (2014, September 18). Endophyte Infection of Tall Fescue and the Impact of Climatic Factors on Ergovaline Concentrations in Field Crops Cultivated in Southern France (world) [Research-article]. ACS Publications; American Chemical Society. 10.1021/jf503015m

49. Richard, A.-M., Gervais, R., Tremblay, G. F., Bélanger, G., & Charbonneau, É. (2020). Tall fescue as an alternative to timothy fed with or without alfalfa to dairy cows. Journal of Dairy Science, 103(9), 8062–8073. 10.3168/jds.2019-18120

50. Roberts, C. A., Andrae, J. G., Smith, S. R., Poore, M. H., Young, C. A., Hancock, D. W., & Pent, G. J. (2020). The Alliance for Grassland Renewal: A Model for Teaching Endophyte Technology. World Academy of Science, Engineering and Technology International Journal of Animal and Veterinary Sciences, 14. https://publications.waset.org/10011192/the-alliance-for-grassland-renewal-a-model-for-teaching-endophyte-technology

51. Schardl, C. L., Grossman, R. B., Nagabhyru, P., Faulkner, J. R., & Mallik, U. P. (2007). Loline alkaloids: Currencies of mutualism. Phytochemistry, 68(7), 980–996. 10.1016/j.phytochem.2007.01.010

52. Schardl, C. L., Panaccione, D. G., & Tudzynski, P. (2006). Ergot alkaloids—Biology and molecular biology. The Alkaloids. Chemistry and Biology, 63, 45–86. 10.1016/s1099-4831(06)63002-2

53. Schnitzius, J. M., Hill, N. S., Thompson, C. S., & Craig, A. M. (2001). Semiquantitative determination of ergot alkaloids in seed, straw, and digesta samples using a competitive enzyme-linked immunosorbent assay. *Journal of Veterinary Diagnostic Investigation: Official Publication of the American Association of Veterinary Laboratory Diagnosticians*, Inc, 13(3), 230–237. 10.1177/104063870101300307

54. Shang, L., He, W., Wang, T., Yang, Y., Xu, Q., Zhao, X., Yang, L., Zhang, H., Li, X., Lv, Y., Chen, W., Cao, S., Wang, X., Zhang, B., Liu, X., Yu, X., He, H., Wei, H., Leng, Y., … Qian, Q. (2023). A complete assembly of the rice Nipponbare reference genome. Molecular Plant, 16(8), 1232–1236. 10.1016/j.molp.2023.08.003

55. Takach, J. E., & Young, C. A. (2014). Alkaloid Genotype Diversity of Tall Fescue Endophytes. Crop Science, 54(2), 667–678. 10.2135/cropsci2013.06.0423

56. Talamantes, D., Kirkpatrick, C., & Wallace, J. (2024). Developing robust quantitative PCR primers for comparative biomass analysis of Tall Fescue (Festuca arundinacea) and its Epichloë endophyte. microPublication Biology. 10.17912/micropub.biology.001275

57. West, C. P. (2007). Plant Influences on Endophyte Expression. 117–121.

58. Wu, T. D., & Nacu, S. (2010). Fast and SNP-tolerant detection of complex variants and splicing in short reads. *Bioinformatics (Oxford*, England), 26(7), 873–881. 10.1093/bioinformatics/btq057

59. Xu, L., Li, X., Han, L., Li, D., & Song, G. (2017). Epichloe endophyte infection improved drought and heat tolerance of tall fescue through altered antioxidant enzyme activity. European Journal of Horticultural Science, 82, 90–97. 10.17660/eJHS.2017/82.2.4

60. Young, C. A., Bryant, M. K., Christensen, M. J., Tapper, B. A., Bryan, G. T., & Scott, B. (2005). Molecular cloning and genetic analysis of a symbiosis-expressed gene cluster for lolitrem biosynthesis from a mutualistic endophyte of perennial ryegrass. Molecular Genetics and Genomics: MGG, 274(1), 13–29. 10.1007/s00438-005-1130-0

61. Young, C. A., Schardl, C. L., Panaccione, D. G., Florea, S., Takach, J. E., Charlton, N. D., Moore, N., Webb, J. S., & Jaromczyk, J. (2015). Genetics, genomics and evolution of ergot alkaloid diversity. Toxins, 7(4), 1273–1302. 10.3390/toxins7041273

62. Zhang, W., Card, S. D., Mace, W. J., Christensen, M. J., McGill, C. R., & Matthew, C. (2017). Defining the pathways of symbiotic Epichloë colonization in grass embryos with confocal microscopy. Mycologia, 109(1), 153–161. 10.1080/00275514.2016.1277469

